# Context dependency of time-based event-related expectations for different modalities

**DOI:** 10.1101/2021.03.06.434208

**Authors:** Felix Ball, Julia Andreca, Toemme Noesselt

## Abstract

Expectations about the temporal occurrence of events (when) are often tied with the expectations about certain event-related properties (what and where) happening at these time points. For instance, slowly waking up in the morning we expect our alarm clock to go off; however, the longer we do not hear it the more likely we already missed it. However, most current evidence for complex time-based event-related expectations (TBEEs) is based on the visual modality. Here we tested whether TBEEs can also act cross-modally. To this end, visual and auditory stimulus streams were presented which contained early and late targets embedded among distractors (to maximise temporal target uncertainty). Foreperiod-modality-contingencies were manipulated run-wise so that visual targets either occurred early in 80% of trials and auditory targets occurred late in 80 % of trials or vice versa. Participants showed increased sensitivity for expected auditory early/visual late targets which increased over time while the opposite pattern was observed for visual early/auditory late targets. A benefit in reaction times was only found for auditory early trials. Together, this pattern of results suggests that context-dependent TBEEs for auditory targets after short foreperiods (be they correct or not) dominated and determined which modality became more expected at the late position irrespective of the veridical statistical regularity. Hence, TBEEs in cross-modal and uncertain environments are context-dependent, shaped by the dominant modality in temporal tasks (i.e. auditory) and only boost performance cross-modally when expectations about the event after the short foreperiod match with the run-wise context (i.e. auditory early/visual late).

## 1. Introduction

Throughout our lives, we learn about regularities and contingencies in our environment and use expectations about them to optimize and adapt our behaviour. For instance, if we go outside, we are likely to encounter potentially harmful cars on the street rather than the sidewalk (event-related expectations) and in sports, the command ‘ready-set-go’ provides temporal cues as to when the start signal will occur (temporal [time-related] expectations). However, often event- and time-related expectations go hand in hand: after putting our favourite dish in the oven, we soon start to expect to smell a delicious scent; however, the longer we do not smell anything the more likely that we forgot to turn on the oven.

In the past, different types of expectations – such as spatial, temporal and identity-specific expectations^1^ – have been examined. Spatial attention and spatial expectations have traditionally been studied by using spatial probabilistic cues, indicating with a certain likelihood where targets will appear (thus, creating expectations about target’s upcoming position) without providing information about the temporal onset or identity of targets (Posner, 1980; Posner et al., 1980; Zuanazzi & Noppeney, 2020). In contrast, time-related or temporal expectations (for review, see Nobre & Rohenkohl, 2014) have been studied by manipulating the likelihood of target’s onset time but here, equally for each event type and spatial position. For instance, target occurrence could be more likely after a short than long interval while the presence of particular stimulus features (which has to be discriminated/identified) is unpredictable (Ball, Michels, et al., 2018; Ball, Fuehrmann, et al., 2018; Ball, Nentwich, et al., 2021; Ball, Spuerck, et al., 2021). Finally, identity-specific expectations – expectations about the identity of the upcoming target – have been studied in many contexts, e.g. by manipulating the likelihood that a specific feature (which has to be discriminated) will be presented, while controlling for target’s temporal onset and spatial occurrence (Puri & Wojciulik, 2008; Summerfield & Egner, 2016). All aforementioned expectation types have been linked to improvements in accuracy and/or reaction times for expected compared to unexpected events and points in time; further, they are mainly studied in the visual domain. While identity-specific expectations are solely based on what will happen, spatial expectations are solely based on where something will happen and temporal expectations (TE) are solely based on when something will happen; in contrast, time-based event-related expectations (TBEE) are expectations for a certain target property of the event (what or where) conditioned upon certain points in time (for review, see Thomaschke & Dreisbach, 2015). Thus, they entail when something will happen but also what or where it will happen.

Like other research on expectation, previous studies on TBEEs were mainly restricted to the visual domain (Aufschnaiter, Kiesel, & Thomaschke, 2018; Aufschnaiter, Kiesel, Dreisbach, et al., 2018; Kunchulia et al., 2017; Thomaschke et al., 2011; Thomaschke & Dreisbach, 2013, 2015; Volberg & Thomaschke, 2017; Wagener & Hoffmann, 2010). In the majority of these studies, TBEEs were manipulated by adjusting the event-foreperiod contingencies; to this end, researchers used two different foreperiods (e.g. 600 ms and 1400 ms, indicated by the presentation of a fixation cross) and rendering one of two possible events (e.g. square or circle) more likely to be presented after one than the other foreperiod (e.g. square appears 80 % after 600 ms and 20 % after 1400 ms and for circle it is the other way around). Note that the number of trials of each foreperiod and event were balanced in these studies, so that only the event-foreperiod contingencies were manipulated. The central finding across studies was that reaction times (but not necessarily accuracy) were reduced for likely compared to unlikely event-foreperiod contingencies. Moreover, TBEEs are found in different contexts similarly to TEs which are observed for different tasks, for different modalities etc. (Ball, Michels, et al., 2020; Ball, Fuehrmann, et al., 2018; Ball, et al., 2020; Ball, Nentwich, et al., 2021; Ball, Spuerck, et al., 2021; Coull & Nobre, 1998; Cravo et al., 2013; Jepma et al., 2012; Niemi & Näätänen, 1981). For instance, reaction times in TBEE studies do not only improve when the foreperiod primes a specific shape that has to be identified (Thomaschke et al., 2011; Wagener & Hoffmann, 2010), but also when it primes words which have to be discriminated (Thomaschke et al., 2018), a specific task that has to be executed (Aufschnaiter, Kiesel, & Thomaschke, 2018; Aufschnaiter, Kiesel, Dreisbach, et al., 2018) or a specific spatial position instead of the to-be-distinguished item itself (Wagener & Hoffmann, 2010). Further, behavioural benefits due to TBEE can extend to time points adjacent to the most likely foreperiod (Thomaschke et al., 2011). This is in close resemblance to findings showing that benefits due to TE can also generalize across larger time windows (see e.g. Ball, Michels, et al., 2018; Bouwer & Honing, 2015; Breska & Deouell, 2016). In sum, previous research demonstrated that TBEEs can act within the visual domain and effectively prime simple visual features, spatial positions as well as more abstract constructs such as the potentially upcoming task itself.

However, TBEEs – so far – have been dominated by investigations in unisensory visual events. As our surrounding often stimulates multiple senses, we here addressed the important question whether TBEEs generalise across different sensory systems and can also be created for stimuli of different modalities. To our knowledge, it is currently unresolved whether TBEEs are operating only within one sensory system or can act in a cross-modal, generalised fashion and thus, prime the appearance of a modality-specific event depending on the foreperiod. Recently, we were able to show that TEs (faster and more correct responses for stimuli presented at expected moments in time) differ across modalities (Ball, Michels, et al., 2018; Ball, Fuehrmann, et al., 2018; Ball, Nentwich, et al., 2021; Ball, Spuerck, et al., 2021); specifically, effects of TE were often reduced for unimodal auditory and even more for visual unisensory targets (compared to multisensory audio-visual targets). This finding might imply that participants, instead of creating TEs – which should facilitate perception of and preparation for all targets independent of modality – may have rather created modality-specific TBEE in our TE study. However, given that we manipulated only TEs and not TBEE, our results can also be explained by multisensory interplay (i.e. higher performance for multi- compared to unisensory stimuli; see e.g. Driver & Noesselt, 2008; Starke et al., 2020) and by the notion that some targets (i.e. audio-visual and auditory) might be easier affected by TEs (as compared to visual targets; see Ball, Fuehrmann, et al., 2018; Ball, Michels, et al., 2018; Wilsch et al., 2020). In contrast to our multisensory TE experiment, three previous studies (Lange & Röder, 2006; Mühlberg et al., 2014; Mühlberg & Soto-Faraco, 2019) used a multisensory hybrid-design (audio-tactile or visual-tactile experiments) in which the authors manipulated TBEEs (one modality more likely depending on time point) but also TEs (one time point was more likely) as well as identity-specific expectation (one modality was more likely)^2^. However, these experiments also do not allow for any strong conclusions about cross-modal TBEEs as either TBEE effects were not analysed (Mühlberg et al., 2014; Mühlberg & Soto-Faraco, 2019) or data analysis was restricted to short foreperiods (Lange & Röder, 2006), i.e. only partially analysed^3^. Further, it is impossible to determine which type of expectation (temporal, identity-specific and/or TBEE) shaped the presented data patterns. For instance, it is conceivable that the mere combination of identity-specific and temporal expectations affected participants’ performance. If TBEEs were affecting performance in these studies, the presented descriptive statistics suggest that they potentially decrease reaction times for the more likely (primary) compared to the less likely presented modality (secondary) but only when presented after an expected foreperiod. Hence, while it is possible that TBEEs could prime a certain modality based on the foreperiod and thereby affecting behaviour, previous studies – due to the lack of focus on this topic – fell short to provide evidence of the existence of cross-modal TBEEs.

Here we investigate for the first time directly whether participants can create TBEEs for events of different modalities. To this end, we altered an audio-visual paradigm which we previously established to study TEs (Ball, Fuehrmann, et al., 2018; Ball, Michels, et al., 2018; Jaramillo & Zador, 2011) based on standard designs to investigate TBEEs (Thomaschke & Dreisbach, 2015; Volberg & Thomaschke, 2017; Wagener & Hoffmann, 2010). Note that our paradigm was specifically designed to test for changes in perceptual sensitivity and response times while previous studies focused mainly on RT effects. In the current study, we balanced the number of events (i.e. modalities) and foreperiod intervals and exclusively manipulated TBEEs. On each trial, we presented a sequence of 15 auditory or visual stimuli (50:50) with one stimulus being the deviant target stimulus that had to be identified (lower or higher frequency than distractors). Based on previous TBEE studies (Thomaschke & Dreisbach, 2015; Volberg & Thomaschke, 2017; Wagener & Hoffmann, 2010), target stimuli were either presented early or late in the sequence (50:50) and TBEEs were manipulated run-wise. In one experimental run, auditory target stimuli were more likely to be presented early (80 %) while visual target stimuli were more likely (80 %) to be presented late. In the other run, the foreperiod-event contingency was reversed. We hypothesized that if TBEEs generally affect behaviour, we should observe faster and more accurate responses to targets which are expected dependent on the specific foreperiod (e.g. auditory instead of visual targets and vice versa). In addition, however, TBEEs might also be context-dependent. Since the auditory modality is better suited for information extraction in temporal contexts (Ball, Fuehrmann, et al., 2018; Ball, Michels, et al., 2018; Bertelson & Aschersleben, 2003; Kemény & Lukács, 2019; Repp & Penel, 2002; Welch et al., 1986; Welch & Warren, 1986; Wilsch et al., 2020), we further hypothesised that TBEEs might be more pronounced for auditory stimuli.

## 2. Methods

### 2.1. Participants

We collected data from 31 participants. All participants provided written informed consent and declared to be free of neurological or psychiatric disorders and to have normal or corrected visual acuity. One participant was excluded due to low performance (performance in at least one condition < 25 % correct; Ball, Fuehrmann, et al. 2018; Ball, Michels, et al. 2018 for identical criteria), leaving 30 participants for analysis (mean age ± SD: 23.6 ± 4.3, # women: 18, # left-handed: 1). This study was approved by the local ethics committee of the Otto-von-Guericke-University, Magdeburg.

### 2.2. Apparatus

The experiments were programmed using the Psychophysics Toolbox (Brainard, 1997) and Matlab 2012b (Mathworks Inc.). Stimuli were presented on a LCD screen (22’’ 120 Hz, SAMSUNG 2233RZ) with optimal timing and luminance accuracy for vision researches (Wang & Nikolić, 2011). Resolution was set to 1650 x 1080 pixels and the refresh rate to 60 Hz. Participants were seated in front of the monitor at a distance of 102 cm (eyes to fixation point). Responses were collected with a wireless mouse (Logitech M325). Accurate timing of stimuli (<= 1 ms) was confirmed prior to the experiment with a BioSemi Active-Two EEG amplifier system connected with a microphone and photodiode.

### 2.3. Stimuli

Stimulus sequences on each trial consisted either of auditory (pure tones) or visual stimuli (circles filled with chequerboards). Chequerboards subtended 3.07° visual angle, were presented above the fixation cross (centre to centre distance of 2.31°) and on a dark grey background (RGB: [25.5 25.5 25.5]). The fixation cross (white) was presented 2.9° above the screen’s centre. As in our previous reports (Ball, Fuehrmann, et al., 2018; Ball, Michels, et al., 2018), sounds were presented from one speaker placed on top of the screen at a distance of 7.06° from fixation, 4.76° from chequerboard’s centre, and 3.22° from chequerboard’s edge. Chequerboards and pure sounds were used as targets and distractors. The distractor frequencies were jittered randomly between 4.6, 4.9, and 5.2 cycles per degree for chequerboards and between 2975, 3000 and 3025 Hz for sounds. Visual and auditory target frequencies were individually adjusted to a 75 % accuracy level at the beginning of the experiment. Hence, targets – although the same type of stimulus (chequerboard/pure sound) – were either lower or higher in frequency compared to distractor frequencies. Furthermore, the intensities for both target and distractor chequerboards and sounds were varied randomly throughout the stimulus sequences. The non-white checkers were jittered between 63.75, 76.5, and 89.25 RGB (average grey value of 76.5 RGB). The sound intensities were jittered between 20 %, 25 %, and 30 % of the maximum sound intensity (average of 25 % = 52 dB[A]).

### 2.4. Procedure

Participants were seated in a dark, sound-attenuated chamber. Each experiment consisted of three parts; first, participants completed one or two training runs (32 trails per run) to familiarize themselves with the task. Next, they completed two threshold determination runs during which the frequency of the sound and chequerboard target stimuli was adjusted to 75 % correct responses. Finally, participants completed 6 experimental runs (160 trials per run, 960 trials total). After completion of the experiment participant were interviewed whether they realised the foreperiod-modality contingencies. The whole procedure (including instructions etc.) took approximately 2.5 − 3 h per participant.

Each trial consisted of a stimulus sequence of 15 stimuli (50 ms stimulus and 100 ms gap), followed by a response window (1500 ms) and an inter-stimulus-interval (350 – 1350 ms) in which no response was recorded. The stimulus sequence of each trial was either auditory or visual and participants were informed that on each trial a target (lower or higher frequency than distractors) was embedded in the sequence (see **Figure 1: Schematic examples for stimulus sequences on each trial (top) and context-dependent probabilities of target stimuli (bottom)**. (Top) Each trial was preceded by a variable Inter-Trial-Interval (ITI, i.e. blank screen). The blank screen (350 - 1350 ms) was followed by a sequence of 15 stimuli (either auditory [see left stimulus sequence] or visual [see right stimulus sequence]). Stimuli were presented for 50 ms with a gap of 100 ms in between successive stimuli. Target stimuli (adjusted to a threshold of 75 % accuracy) are highlighted in this example with a red contour (not present in the experiment). Target stimuli were either higher or lower in frequency as compared to distractors and were presented either early (3^rd^ position) or late (11^th^, 13^th^ or 15^th^ position) in the sequence. After the stimulus sequence ended, participants had another 1500 ms for providing their response. (Bottom) Depiction of the context-dependent probabilities of modality, foreperiod and modality-foreperiod contingencies in each run. In both run types (auditory early [upper row] and visual early runs [lower row]), there was an equal probability (50 %) for each modality and foreperiod to be presented (left and middle plots). Only the modality-foreperiod contingencies (right plots) were altered (80 % vs. 20 %) across runs to exclusively modulate time-based expectations., top). Participants were asked to discriminate the frequency of each target as quickly and accurately as possible. Participants held the response device (i.e. mouse) with both hands, while placing their left/right thumbs on the left/right mouse buttons, respectively. Each button was used for one of the two response options (low/high frequency; key bindings were counterbalanced across participants). The response recording started with the onset of the first stimulus of the sequence and ended 1500 ms after sequence’s offset (response window). Only the first button press was recorded. In case no button was pressed, the trial was repeated at the end of each run’s quarter (mean of repeated trials across participants: 2.2 ± 2.7 % SD).

**Figure 1:**
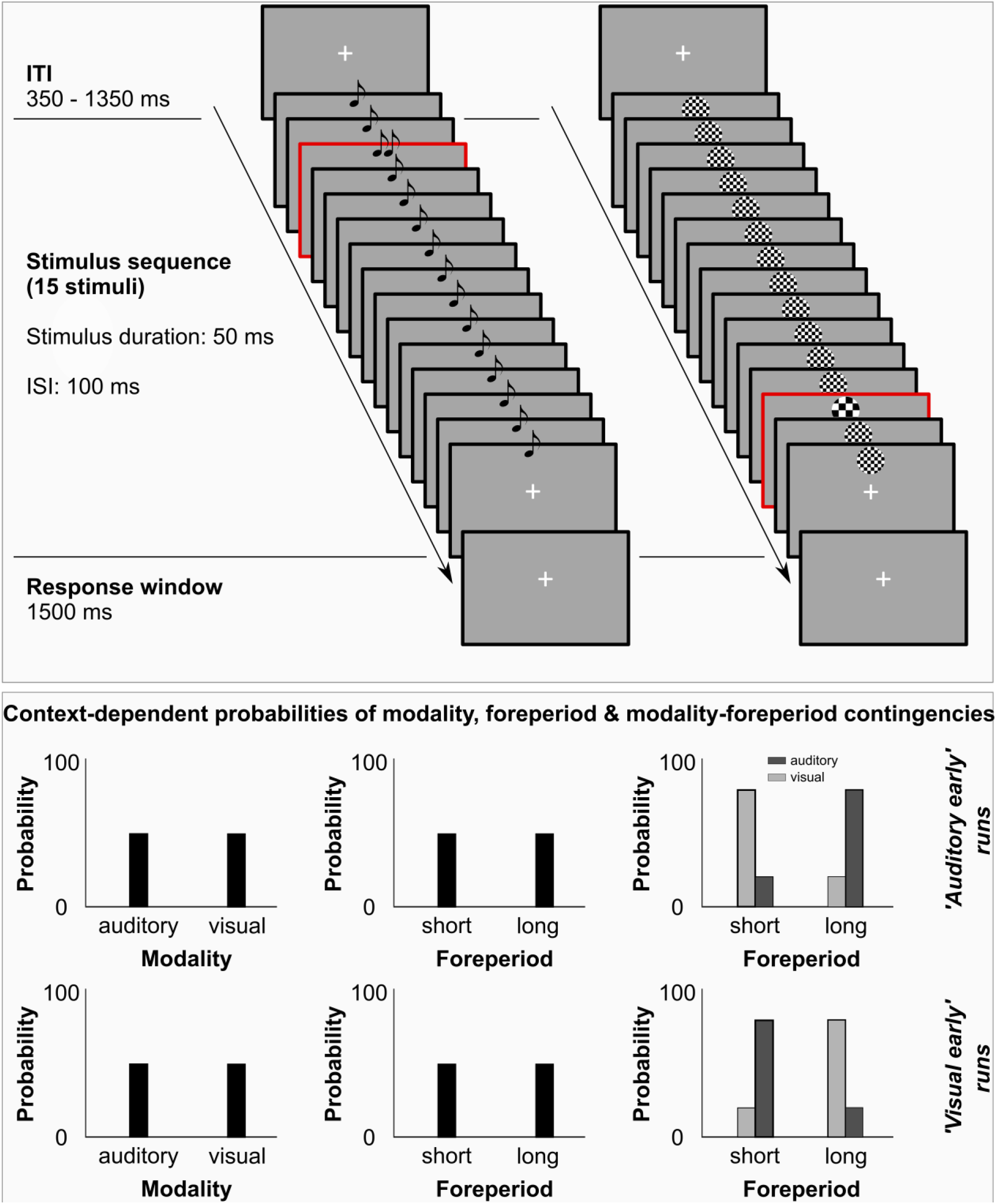
Schematic examples for stimulus sequences on each trial (top) and context-dependent probabilities of target stimuli (bottom). (Top) Each trial was preceded by a variable Inter-Trial-Interval (ITI, i.e. blank screen). The blank screen (350 - 1350 ms) was followed by a sequence of 15 stimuli (either auditory [see left stimulus sequence] or visual [see right stimulus sequence]). Stimuli were presented for 50 ms with a gap of 100 ms in between successive stimuli. Target stimuli (adjusted to a threshold of 75 % accuracy) are highlighted in this example with a red contour (not present in the experiment). Target stimuli were either higher or lower in frequency as compared to distractors and were presented either early (3^rd^ position) or late (11^th^, 13^th^ or 15^th^ position) in the sequence. After the stimulus sequence ended, participants had another 1500 ms for providing their response. (Bottom) Depiction of the context-dependent probabilities of modality, foreperiod and modality-foreperiod contingencies in each run. In both run types (auditory early [upper row] and visual early runs [lower row]), there was an equal probability (50 %) for each modality and foreperiod to be presented (left and middle plots). Only the modality-foreperiod contingencies (right plots) were altered (80 % vs. 20 %) across runs to exclusively modulate time-based expectations.

Within each run, we balanced the number of low and high frequency targets, auditory and visual sequences, as well as the number of targets appearing after a short (early position: 3^rd^ position [50 % of trials]) or long foreperiod (late positions [50 % of trials]). Thus overall, each of these events were equally likely (see **Figure 1: Schematic examples for stimulus sequences on each trial (top) and context-dependent probabilities of target stimuli (bottom). (Top) Each trial was preceded** by a variable Inter-Trial-Interval (ITI, i.e. blank screen). The blank screen (350 - 1350 ms) was followed by a sequence of 15 stimuli (either auditory [see left stimulus sequence] or visual [see right stimulus sequence]). Stimuli were presented for 50 ms with a gap of 100 ms in between successive stimuli. Target stimuli (adjusted to a threshold of 75 % accuracy) are highlighted in this example with a red contour (not present in the experiment). Target stimuli were either higher or lower in frequency as compared to distractors and were presented either early (3^rd^ position) or late (11^th^, 13^th^ or 15^th^ position) in the sequence. After the stimulus sequence ended, participants had another 1500 ms for providing their response. (Bottom) Depiction of the context-dependent probabilities of modality, foreperiod and modality-foreperiod contingencies in each run. In both run types (auditory early [upper row] and visual early runs [lower row]), there was an equal probability (50 %) for each modality and foreperiod to be presented (left and middle plots). Only the modality-foreperiod contingencies (right plots) were altered (80 % vs. 20 %) across runs to exclusively modulate time-based expectations., bottom left and middle column). However, to modulate TBEEs, we manipulated the ‘foreperiod-event’ contingencies and thus, the context within each run. In ‘auditory early’ runs, auditory target stimuli were more likely (80 %) to appear at the early position, while visual stimuli were more likely to appear (80 %) at the late position. In ‘visual early’ runs, these likelihoods were reversed (see **Figure 1: Schematic examples for stimulus sequences on each trial (top) and context-dependent probabilities of target stimuli (bottom).**(Top) Each trial was preceded by a variable Inter-Trial-Interval (ITI, i.e. blank screen). The blank screen (350 - 1350 ms) was followed by a sequence of 15 stimuli (either auditory [see left stimulus sequence] or visual [see right stimulus sequence]). Stimuli were presented for 50 ms with a gap of 100 ms in between successive stimuli. Target stimuli (adjusted to a threshold of 75 % accuracy) are highlighted in this example with a red contour (not present in the experiment). Target stimuli were either higher or lower in frequency as compared to distractors and were presented either early (3^rd^ position) or late (11^th^, 13^th^ or 15^th^ position) in the sequence. After the stimulus sequence ended, participants had another 1500 ms for providing their response. (Bottom) Depiction of the context-dependent probabilities of modality, foreperiod and modality-foreperiod contingencies in each run. In both run types (auditory early [upper row] and visual early runs [lower row]), there was an equal probability (50 %) for each modality and foreperiod to be presented (left and middle plots). Only the modality-foreperiod contingencies (right plots) were altered (80 % vs. 20 %) across runs to exclusively modulate time-based expectations., bottom right column). Note that our previous work showed that basic TEs for early and late positions differ (Ball, Fuehrmann, et al., 2018; Ball, Michels, et al., 2018); as long as the target stimulus is not presented, expectation exponentially increases that the target will soon be presented (hazard rate). Thus, accuracy is e.g. higher for the late position. To counter the hazard rate effect, we used different late positions (11^th^ [31.25 %], 13^th^ [12.5 %] or 15^th^ [6.25 %]) to render expectations for the early and late positions more alike.

### 2.5. Data analyses

For analyses, we used Matlab 2017b (Mathworks Inc.) and JASP (v 13.0.0). To increase comparability across studies, we applied the most common outlier exclusion criteria as used in previous studies on TBEEs (Aufschnaiter, Kiesel, & Thomaschke, 2018; Aufschnaiter, Kiesel, Dreisbach, et al., 2018; Kunchulia et al., 2017; Thomaschke et al., 2011; Thomaschke & Dreisbach, 2013, 2015; Volberg & Thomaschke, 2017; Wagener & Hoffmann, 2010): for each condition we excluded RTs below and above 3 times the standard deviation around the mean and response times below 100 ms, resulting in an average exclusion of 2.3 ± 2.9 % of trials per condition.

To decrease familywise error rates of the analysis and since context dependency of modality-related TBEEs was the main focus of this investigation, we calculated the difference in performance between the primary (i.e. expected, more likely) and secondary (i.e. unexpected, less likely) condition and used this difference for statistical analyses (Eimer & Kiss, 2008; Luck & Gaspelin, 2017; Näätänen et al., 2004; Sawaki et al., 2012; Thomas & Zumbo, 2012; Zimmerman et al., 1993). This difference was calculated for each *foreperiod* (short, long), *context* (auditory early [= higher likelihood for auditory early and visual late], visual early [= higher likelihood for visual early and auditory late]) and *run half* (first, second) which also constituted the within-subject factors of the analysis. We included the factor *run half* to test whether TBEE improves over time as participants have to re-learn the time-based regularities in each run. The factor *contex*t was included as participants might learn one time-modality contingency (e.g. auditory early and visual late) but not the other. Finally, we also added the between-subject factor *first run* (‘auditory early’, ‘visual early’) to account for potential interaction effects with factor *context*, i.e. a bias introduced by the likelihoods the participants encountered first. We conducted two repeated-measures ANOVAs, one for accuracy and one for response times. If required, ANOVA results were Greenhouse-Geisser (p_GG_) corrected. Further, we used the ‘simple main effects’ function in JASP for post-hoc tests.

## 3. Results

Our dependent variable, the difference between the primary (expected) and secondary (unexpected) condition, indicating the presence of TBEEs, was calculated so that positive values always indicate higher performance in the primary condition (higher accuracy and faster reaction times) to render results of both measures more easily comparable.

The results of the repeated measures ANOVA support the notion that modality-related TBEEs are context-dependent. For differences in accuracy there was a trend for an effect of *context* (auditory early run vs. visual early run: F(1,28) = 3.681, p = .065, η_p_^2^ = .116) which was significantly interacting with the factor *run half* (*context * run half*: F(1,28) = 6.166, p = .019, η_p_^2^ = .18). TBEE effects increased over time in ‘auditory early’ runs (F = 4.49, p = .043) and – if at all – decreased over time in ‘visual early’ runs (F = 2.568, p = .12) as can be seen in Figure 2: Accuracy and response time results for significant interaction effects. (A) Interaction effect of context and run half for accuracies. (B) Descriptive statistics of all factor levels (foreperiod, run half and context) for differences in accuracy. (C and D) Same descriptive statistics as A and B for response times. Note that we plot the TBEE effect – hence, the difference between the primary and secondary condition. Positive values always indicate higher performance (higher accuracy or faster response times) in the primary compared to the secondary condition. Error bars are standard errors of the mean. A. All remaining effects were non-significant (all F < 2.596, all p >.118; for effects specifically including factor *first run*: all p > .184). Nevertheless, we present group-mean averages from all conditions in Figure 2: Accuracy and response time results for significant interaction effects. (A) Interaction effect of context and run half for accuracies. (B) Descriptive statistics of all factor levels (foreperiod, run half and context) for differences in accuracy. (C and D) Same descriptive statistics as A and B for response times. Note that we plot the TBEE effect – hence, the difference between the primary and secondary condition. Positive values always indicate higher performance (higher accuracy or faster response times) in the primary compared to the secondary condition. Error bars are standard errors of the mean. B for better comparison with the response time results (see below and Figure 2: Accuracy and response time results for significant interaction effects. (A) Interaction effect of context and run half for accuracies. (B) Descriptive statistics of all factor levels (foreperiod, run half and context) for differences in accuracy. (C and D) Same descriptive statistics as A and B for response times. Note that we plot the TBEE effect – hence, the difference between the primary and secondary condition. Positive values always indicate higher performance (higher accuracy or faster response times) in the primary compared to the secondary condition. Error bars are standard errors of the mean. C & D for RT results).

**Figure 2:**
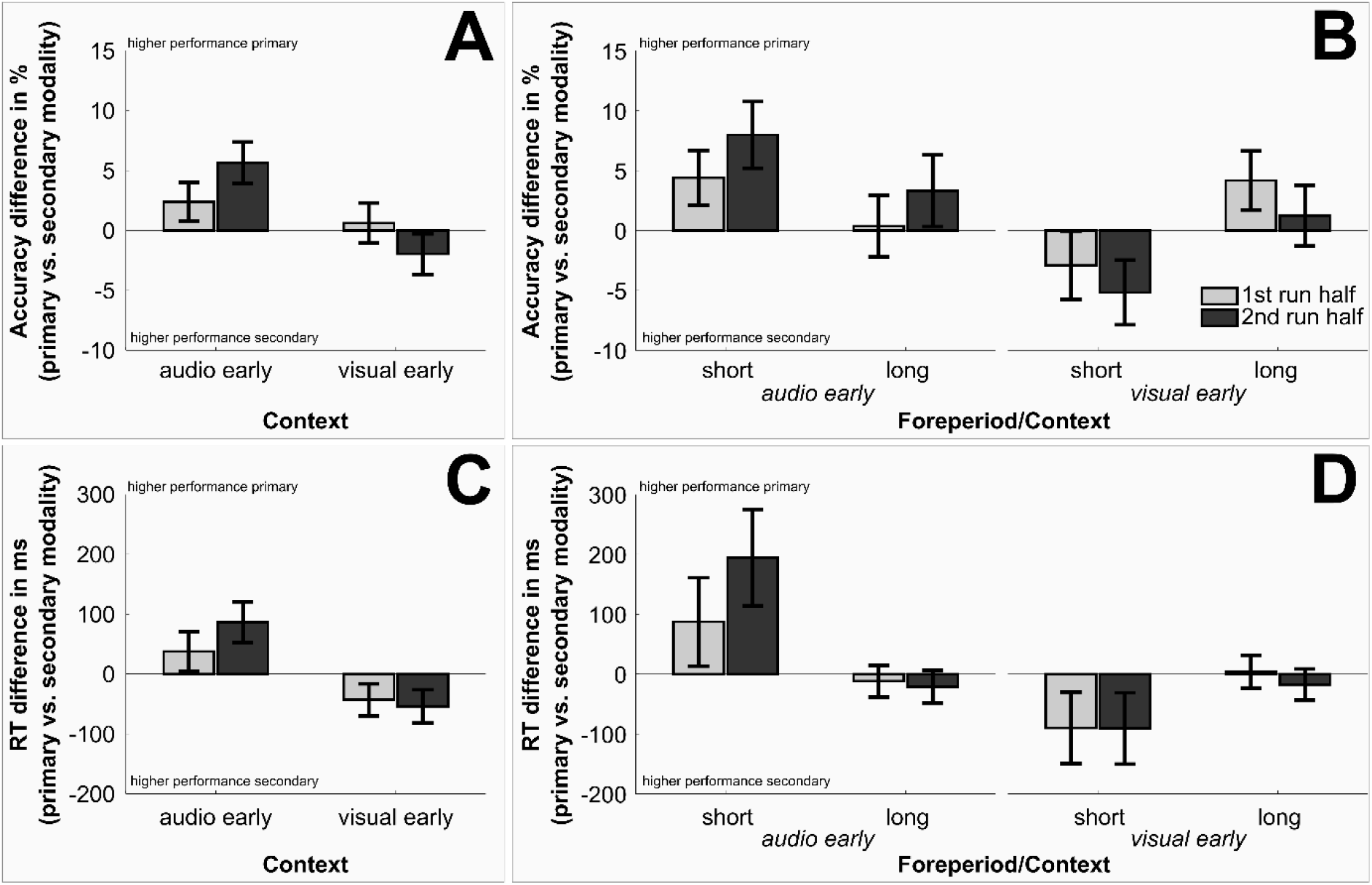
Accuracy and response time results for significant interaction effects. (A) Interaction effect of context and run half for accuracies. (B) Descriptive statistics of all factor levels (foreperiod, run half and context) for differences in accuracy. (C and D) Same descriptive statistics as A and B for response times. Note that we plot the TBEE effect – hence, the difference between the primary and secondary condition. Positive values always indicate higher performance (higher accuracy or faster response times) in the primary compared to the secondary condition. Error bars are standard errors of the mean. Bright grey bars indicate data of the first half and dark grey bars the second half of the run.

For response times we observed corresponding results. Again, there was a trend for an effect of *context* (F(1,28) = 4.112, p = .052, η_p_^2^ = .128) which was once more significantly interacting with the factor *run half* (context * run half: F(1,28) = 4.276, p = .048, η_p_^2^ = .132). As for accuracies, TBEE increased significantly over time in ‘auditory early’ runs (F = 6.185, p = .019), while this effect was non-significant in ‘visual early’ runs (F = .185, p = .67) as can be seen in Figure 2: Accuracy and response time results for significant interaction effects. (A) Interaction effect of context and run half for accuracies. (B) Descriptive statistics of all factor levels (foreperiod, run half and context) for differences in accuracy. (C and D) Same descriptive statistics as A and B for response times. Note that we plot the TBEE effect – hence, the difference between the primary and secondary condition. Positive values always indicate higher performance (higher accuracy or faster response times) in the primary compared to the secondary condition. Error bars are standard errors of the mean. C. Further, there was a trend (F(1,28) = 3.452, p = .074, η_p_^2^ = .11) for an interaction of factors *first run* and *context*: TBEEs were increased in ‘auditory early’ runs when participants started with the ‘auditory early’ as compared to ‘visual early’ runs (avg. diff = 122 ms, F = 3.894, p = .058). In ‘visual early’ runs, TBEE effects rather decreased and inverted (although non-significantly) when participants started with the ‘auditory early’ as compared to ‘visual early’ runs (avg. diff = − 70 ms, F = 1.941, p = .175). Finally, the triple interaction of factors context, run half and foreperiod (see Figure 2: Accuracy and response time results for significant interaction effects. (A) Interaction effect of context and run half for accuracies. (B) Descriptive statistics of all factor levels (foreperiod, run half and context) for differences in accuracy. (C and D) Same descriptive statistics as A and B for response times. Note that we plot the TBEE effect – hence, the difference between the primary and secondary condition. Positive values always indicate higher performance (higher accuracy or faster response times) in the primary compared to the secondary condition. Error bars are standard errors of the mean. D) was significant (F(1,28) = 4.183, p = .05, η_p_^2^ = .13), indicating that the increase in TBEE over time was mainly driven by the short foreperiod trials in ‘auditory early’ runs (F = 6.815, p = .014) but less by all other conditions (all F < .964, all p > .335). All remaining effects were non-significant (all F < 2.847, all p >.103; for effects specifically including factor *first run*: all p > .125).

After the experiment, none of the participants reported to have noticed that we manipulated foreperiod-modality contingencies within and across runs. This held true, even after being informed about the experimental manipulation.

Finally, we would like to direct readers’ attention to the general pattern of accuracies and response times in Figure 2: Accuracy and response time results for significant interaction effects. (A) Interaction effect of context and run half for accuracies. (B) Descriptive statistics of all factor levels (foreperiod, run half and context) for differences in accuracy. (C and D) Same descriptive statistics as A and B for response times. Note that we plot the TBEE effect – hence, the difference between the primary and secondary condition. Positive values always indicate higher performance (higher accuracy or faster response times) in the primary compared to the secondary condition. Error bars are standard errors of the mean. C and Figure 2: Accuracy and response time results for significant interaction effects. (A) Interaction effect of context and run half for accuracies. (B) Descriptive statistics of all factor levels (foreperiod, run half and context) for differences in accuracy. (C and D) Same descriptive statistics as A and B for response times. Note that we plot the TBEE effect – hence, the difference between the primary and secondary condition. Positive values always indicate higher performance (higher accuracy or faster response times) in the primary compared to the secondary condition. Error bars are standard errors of the mean. D. Depending on the context (auditory or visual early), accuracy-related time-based attention appears to either increase (auditory early) or decrease (visual early) over run half. Generally, expectation effects appear to be reversed in ‘visual early’ runs, indicating that performance was higher for the secondary condition. More importantly, even if TBEEs exist in the beginning of the ‘visual early’ runs (see long foreperiod), participants later (2^nd^ run half) focus on the secondary modality at each foreperiod which is also evident for response times.

## 4. Discussion

Here we tested whether time-based event-related expectations (TBEEs), expectations for a certain target stimulus (here a certain modality) contingent upon a specific temporal foreperiod, are generalizable to cross-modal contexts. We found that modality-related TBEEs were observable but context-dependent. Accuracy and response times improved over the course of a run for the primary (expected, more-likely) compared to the secondary (unexpected, less-likely) target condition but only in ‘auditory early’ runs (80 % likelihood that auditory targets are presented early and visual targets are presented late within a sequence of 15 stimuli). In ‘visual early’ runs, performance rather improved over time for the secondary instead of the primary modality. Finally, response times improvements in ‘auditory early’ runs were mainly driven by targets appearing after a short foreperiod.

Our results show for the first time that humans are able to create TBEEs for stimuli of different modalities, thereby significantly extending previous studies restricted to the visual domain (Aufschnaiter, Kiesel, & Thomaschke, 2018; Aufschnaiter, Kiesel, Dreisbach, et al., 2018; Kunchulia et al., 2017; Thomaschke et al., 2011; Thomaschke & Dreisbach, 2013, 2015; Volberg & Thomaschke, 2017; Wagener & Hoffmann, 2010). Moreover, previous studies were typically designed to measure response time differences (see e.g. Volberg & Thomaschke, 2017), resulting in the use of rather ‘simple’ designs (i.e. the foreperiod was followed by only a single, clearly visible stimulus such as a square or circle) while accuracies were at ceiling. Here, we used a more complex paradigm in which targets were embedded among and had thus to be distinguished from distractors. As a result, we were able to show that TBEEs can affect accuracies – and thus, the perception of events – as well as response times. In addition, we show that modality-related TBEEs do not only affect that a certain modality is expected at a certain point in time, but also that this expectation can influence performance in an orthogonal task (frequency discrimination).

The crucial finding of our study is that participants appear to have unknowingly learned time-modality contingencies but only when the auditory target was more likely presented early and the visual presented late as indexed by performance increases (mainly accuracy differences) for the primary target. As we have already discussed previously (TE; Ball, Fuehrmann, et al., 2018; Ball, Michels, et al., 2018), the auditory system has a higher temporal resolution as compared to the visual system, which might render the auditory system more prone to detect and utilise temporal information (see also Bertelson & Aschersleben, 2003; Kemény & Lukács, 2019; Repp & Penel, 2002; Welch & Warren, 1986). This notion was very recently further supported by an MEG study, showing “that spatial attention has a stronger effect in the visual domain, whereas TE effects are more prominent in the auditory domain” (Wilsch et al., 2020). It is possible that the higher temporal acuity of the auditory system allows for registering early auditory (as compared to visual) targets more easily and reliably. This assumption would be in line with the overall descriptive statistics presented in Figure 2: Accuracy and response time results for significant interaction effects. (A) Interaction effect of context and run half for accuracies. (B) Descriptive statistics of all factor levels (foreperiod, run half and context) for differences in accuracy. (C and D) Same descriptive statistics as A and B for response times. Note that we plot the TBEE effect – hence, the difference between the primary and secondary condition. Positive values always indicate higher performance (higher accuracy or faster response times) in the primary compared to the secondary condition. Error bars are standard errors of the mean. . It appears that after a short foreperiod, expectations about early auditory targets are always stronger because even in the ‘visual early’ runs, the secondary auditory target was expected more strongly. Additionally, expectations for early auditory targets increased in both cases over time (1^st^ vs. 2^nd^ run half). Given that we found no significant influence of the first run (visual or auditory early) on performance, it is more likely that this is a general effect rather than a transition of expectations across runs (which should depend on regularities in the first run). Further, general expectations of early auditory targets appear to result in two opposite effects depending on the run type: over the time course of the run, they boost expectations about late visual targets (in ‘auditory early’ runs) and eliminate – previously correctly established – expectations about late auditory targets (in ‘visual early’ runs). Note again, that we used identical stimuli in both runs and only changed the foreperiod-target modality contingencies. Hence, the differential pattern observed for the two run types indicates that participant’s intrinsic expectations affected performance and when matching with the existing statistical regularities, improved the behavioural outcome.

The specificity and effectiveness of time-based expectations can even be further broken down to the expected early auditory target in ‘auditory early’ runs. This was the only condition for which we found an improvement of accuracies as well as response times. For the expected visual target after the long foreperiod (within the same run), we only found an improvement of accuracies and this improvement was restricted to the second half of the run. Note that this accuracy pattern suggests that participants can learn time-based event regularities for a specific context (i.e. ‘auditory early and visual late’), but that they might also be prone to establish even stronger TBEE for a very specific combination of foreperiod and modality (expected auditory early target). The latter suggestion would be in line with our previous finding of stronger auditory than visual TE effects after short foreperiods (Ball, Fuehrmann, et al., 2018; Ball, Michels, et al., 2018). Further, our current and previous results indicate that the use of visual information – especially related to short foreperiods – in tasks including expectations in the temporal domain (be it pure TE or TBEE tasks) appears to be somehow suppressed by the presence of the more informative auditory modality.

Perception of early expected auditory targets facilitates the creation of TBEE for early auditory targets but also for expected late visual targets within the same run. However, in ‘visual early’ runs, early auditory targets still appear to be prioritized, a bias that strengthens over time and even eliminates correct TBEE for the late auditory target. Thus, participants do not exhibit a bias towards the auditory modality in general (which would have facilitated auditory performance irrespective of the foreperiod); rather, any expectation at the short foreperiod (be it correct or incorrect) appears to strengthen over time and determines which event is expected at the late position. Although, our previous work (Ball et al., 2020; Ball, Spuerck, et al., 2021) indicated that explicit temporal knowledge has rather little impact on performance in temporal tasks, it is possible that explicitly attending a certain modality might affect multisensory TBEE. As mentioned in the introduction, if TBEE was affecting performance in previous studies (Lange & Röder, 2006; Mühlberg et al., 2014; Mühlberg & Soto-Faraco, 2019), their descriptive statistics suggest that it improves performance for the expected compared to the unexpected modality (irrespective of the specific modality) after short foreperiods. Given that participants were forced in these studies to actively attend to a certain modality and time point this might have minimized self-determined ‘modality preferences’, resulting in a stable primary vs. secondary modality and thus, context-independent TBEE effect. However, by using such manipulation of attention, these studies also reduce incidental, statistical learning. At this point, future studies are required to determine potential differences between TBEEs based on different paradigms and explicit vs. implicit learning of regularities.

To close with, previous studies discussed the relation of TEs and TBEEs and their role in response preparation and perceptual facilitation. For instance, Mühlberg & Soto-Faraco (2019) argued that TBEE effects – unlike TE effects – do not depend on additional deployment of temporal attention to specific points in time and are solely explained by motor preparation. First, there is no consensus whether temporal expectation and temporal attention are either the same entity, different mechanisms or whether both – temporal and time-based event-related expectations (created due to statistical regularities) – guide temporal attention to relevant points in time. All possibilities can explain the reported results in the TE and TBEE literature. However, whenever discussed, the ‘expectation driven guidance of temporal attention’ hypothesis appears to be the most prominent one in both research areas (Mühlberg & Soto-Faraco, 2019; Nobre & Rohenkohl, 2014; Nobre & van Ede, 2017; Thomaschke & Dreisbach, 2015; Wagener & Hoffmann, 2010). Given that TEs and TBEEs are mostly studied in implicit task regimes (i.e. implicit statistical learning), this would also imply that attention is shifted implicitly, which is in line with the present results and other proposals of implicit attentional shifts in the temporal, spatial and feature domain (Addleman et al., 2019; Balci & Simen, 2014; Ball et al., 2020; Ball, Spuerck, et al., 2021; Bolger et al., 2013; Melcher et al., 2005; Thomaschke et al., 2011; Thomaschke & Dreisbach, 2015). Hence, it is unclear why TE should guide temporal attention and TBEE should not. Second, note that the argument by Mühlberg and Soto-Faraco is partially derived from the finding that TBEEs generalize to other time points than the most likely foreperiods (Thomaschke et al., 2011). However, if a generalisation of TBEE across a broader time window implies the absence of the guidance of temporal attention to relevant points in time, than this would also cause problems for the interpretation of TE results. The spread of TEs (and thus, the time window in which temporal attention is potentially deployed) is rarely analysed in TE studies and e.g. we, like others, were able to show that TEs can operate in a broader time window instead of being exclusive to one specific point in time (see e.g. Ball, Michels, et al., 2018; Bouwer & Honing, 2015; Breska & Deouell, 2016; Jaramillo & Zador, 2011). Hence, both types of expectations appear to share commonalities, with larger performance effects due to temporal attention at the most likely point in time but also, albeit smaller effects at flanking time points (see e.g. Jones et al., 2002; Thomaschke et al., 2011). Finally, the argument that TBEEs are solely explained by motor preparations while TEs are not, is also debatable. In contrast, here we show that accuracies and response times are affected by TBEEs indicating that not only potential motor preparations were affected but also target perception. Further, most TBEE studies were also not designed to test perceptual effects (as pointed out by the authors) but specifically aimed at inducing response time differences due to TBEEs which were later linked to motor preparation effects (Thomaschke & Dreisbach, 2013; Volberg & Thomaschke, 2017). More importantly, even studies on TEs commonly only modulate and report response time effects (for review, see Nobre & Rohenkohl, 2014). However, when specifically testing whether response time changes in the presence of ceiling accuracy indicate motor preparation or perceptual facilitation effects, previous studies reported to be unable to distinguish between these two concepts (Jepma et al., 2012) or reported that response preparation and strategy shifts (instead of perceptual facilitation) likely caused RT modulations (Ball et al., 2020; van den Brink et al., 2020). These findings again, do not only highlight commonalities between TE and TBEE but also highlight that results have to be interpreted cautiously when only response speed but not accuracy is modulated. Together, the present and earlier results suggest that TEs and TBEEs can not only affect motor preparation but also perceptual processing and that TBEEs are likely event-specific instead of event-unspecific temporal expectations (see e.g. Thomaschke et al., 2011; Thomaschke & Dreisbach, 2015; Volberg & Thomaschke, 2017).

## 5. Conclusion

Our results strongly suggest that TBEEs extend to cross-modal contexts, depend on the creation of event-specific temporal expectations, thereby enhancing performance. This performance enhancement can be linked to perceptual in addition to motor facilitation. Most importantly, our results indicate that TBEEs can be context-dependent and seem to be driven by auditory information, if available, especially after short foreperiods. This has crucial implications for multisensory studies of TEs as participants might develop TBEEs instead of TEs if modalities ‘compete for temporal expectation and attention’ within one experiment. More importantly, the competition which appears to be driven by targets presented after short foreperiods results in modality-specific TBEEs (i.e. auditory early) that only improve behaviour cross-modally when it matches with the global context (i.e. statistical regularities) within one run (i.e. auditory early/visual late).

## Declarations

### CRediT author statement

**Felix Ball:**Conceptualization, Methodology, Formal analysis, Investigation, Writing - Original Draft, Writing - Review & Editing, Visualization, Supervision, Project administration, Funding acquisition **Julia Andreca:**Visualization, Writing - Original Draft, Writing - Review & Editing **Toemme Noesselt:**Conceptualization, Methodology, Writing - Original Draft, Writing - Review & Editing, Supervision, Funding acquisition

### Ethics approval

This study was approved by the local ethics committee of the Otto-von-Guericke-University, Magdeburg.

### Consent to participate and publish

Informed consent was obtained from all individual participants included in the study.

### Open Practices Statement

The data and materials are available in the supplementary information files and upon request (e.g. codes). None of the experiments were preregistered.

### Data availability statement

The authors declare that the analysed data as well as potential supplementary analyses (not reported in the main manuscript) are available in the supplementary information files.

### Code availability

Data were analysed with JASP (freely available) and Matlab 2017b (Mathworks Inc.). Any relevant code is available upon request.

## Footnotes

1 In several studies, Thomaschke and colleagues coined the term ‘time-based event expectation’. Thomaschke defined ‘event expectations’ as expectations about what and/or where something will happen. As one might easily misinterpret ‘event expectations’ as expectations e.g. for a specific object (what), we thought it necessary to first distinguish between three main concepts of expectations: temporal (when), spatial (where) and identity-specific (what) expectations. The latter might still be subdivided (e.g. into feature- and object-based expectations), a discussion which is outside the scope of this paper. Importantly, we use the term ‘event-related expectations’ to imply that these expectations can entail every property of the target – what, where and when – although the latter is already covered by the term ‘time-based’.

2 As an example, in each run (124 trials total), there was an imbalance of possible foreperiods (e.g. 88 trials ‘short’ vs. 36 trials ‘long’), modalities (e.g. 88 trials ‘visual’ vs. 36 trials ‘tactile’) as well as their combinations (e.g. 72 trial ‘visual and short’ etc.).

3 Please note that this is not a direct shortcoming of the studies themselves as all three studies focussed on cross-modal de-/coupling of temporal attention and not time-based event-related expectations.

